# A Unified Genomic Mechanism of Cell-Fate Change

**DOI:** 10.1101/2021.11.24.469848

**Authors:** Masa Tsuchiya, Alessandro Giuliani, Giovanna Zimatore, Jekaterina Erenpreisa, Kenichi Yoshikawa

**Affiliations:** SEIKO Life Science Laboratory, SEIKO Research Institute for Education, Osaka 540-659, Japan; Environment and Health Department, Istituto Superiore di Sanitá, 00161 Rome, Italy; eCampus University, 22060 Novedrate, Como, Italy and CNR-IMM Bologna, Italy; Latvian Biomedical Research & Study Centre, Riga, Latvia; Faculty of Life and Medical Sciences, Doshisha University, Kyotanabe 610-0394, Japan

**Keywords:** Genome Expression, Cell-Fate Decision, Self-Organized Criticality (SOC), Critical Point, Genome-Engine, Genome-Attractor, Biological Statistical Mechanics

## Abstract

The purpose of our studies is to elucidate the nature of massive control of whole genome expression with a particular emphasis on cell-fate change. Whole genome expression is coordinated by the emergence of a critical point (CP: a peculiar set of bi-phasic genes) through the genome-engine. In response to stimuli, the genome expression self-organizes three critical states, each exhibiting distinct collective behaviors with its center of mass acting as a local attractor, coexisting with whole genome attractor (GA). Genome-engine mechanism accounts for local attractors interaction in phase space. The CP acts as the organizing center of cell-fate change, and its activation makes local perturbation spread over the genome affecting GA. The activation of CP is in turn elicited by ‘hot-spots’, genes with elevated temporal variance, normally in charge to keep genome expression at pace with microenvironment fluctuations. When hot-spots oscillation exceeds a given threshold, the CP synchronizes with the GA driving genome expression state transition. The expression synchronization wave invading the entire genome depends on the power law fusion-bursting dynamics of silencing pericentromere-associated heterochromatin domains and the consequent folding-unfolding status of transcribing euchromatin domains. The proposed mechanism is a unified step toward a time-evolutional transition theory of biological regulation.

## 1.1 Introduction: Self-Organization of Genome Expression

One of most remarkable discoveries in bioscience is that ectopic expression of a few transcription factors guides reprogramming directly from a somatic cell back to a type of pluripotent stem cell [Takahashi and Yamanaka 2006; G. Liu at al. 2020]. Such a drastic state change involves the coherent on/off switching of thousands of functionally heterogeneous genes together with epigenetic modifications. This mechanism acts through the complex spatio-temporal self-organization of the genome involving on/off switching of a huge number of genes in a remarkably cooperative manner [Chang et al. 2006; MacArthur et al. 2009].

The existence of master regulators of transcription factor-mediated genes [Ohno 1979] has been demonstrated to guide cell-fate changes, where a cascade of transcription factors occurs to induce or suppress coordinated expression of thousands of genes [Takahashi and Yamanaka 2016]. But to give a physically consistent basis to the above observation, we face two fundamental difficulties:

1. The first one is the lack of a sufficient number of regulatory molecules in a cell, in addition to the target on DNA, to reach a stable thermodynamic state (i.e., breakdown of the mass action law accompanied by the violence of ‘law of large number’). The low copy number of specific gene mRNAs provokes stochastic noise [Ferreiro at al. 2011] thereby inducing a substantial instability of genetic product concentrations falsifying any gene-by-gene feedback control hypothesis [Raser and O’Shea 2005; Yoshikawa 2002a].
2. The second difficulty derives from the huge linear dimension of human DNA molecule (around 2 meters) with respect to cell nucleus that makes chromatin very far from a Turing-like string freely accessible by regulatory molecules at single gene level.

As a matter of fact, we must admit that current understanding on the higher-order structural change in relation to the mechanism of control of several thousand genes still remain in a primitive stage even at the present era.

Since the discovery of induced pluripotent stem cell (iPS cell) in 2006 [Takahashi and Yamanaka 2006], it is still a daunting challenge to elucidate the mechanism of how the transition to a different mode of global gene expression (i.e., reprogramming of genome expression) occurs in the entire genome. A fundamental issue, here is to elucidate the self-organization mechanism at the ‘whole genome’ level of gene-expression regulation. This is responsible for the self-control of on/off switching for thousands of functionally unique heterogeneous genes in a small and highly packed cell nucleus.

In (non-equilibrium) statistical mechanics, self-organized criticality (SOC) was proposed as a general theory of complexity to describe self-organization and emergent order in thermodynamically open systems. With considerable research over the past several decades, SOC has become important in many research domains of natural science such as brain research as well as social science. However, a universal classification has not yet been developed to construct a general mathematical formulation of SOC [Marković and Gros 2014]. Useful background information on SOC is found in [Clar et al. 1996; Jensen 1998; Kinouchi and Prado 1999; Turcotte 2001; Halley and Winkler 2008; Plenz and Niebur 2014; Sanchez and Newman 2015].

Our recent studies [Tsuchiya et al. 2014, 2015, 2016, 2017, 2020a; and Giuliani et al., 2018] demonstrated that self-organized criticality (SOC) is a physically motivated candidate for massive, coordinated gene expression regulation. The main purpose of this review paper is to update a unified genomic mechanism of cell-fate change [Tsuchiya at al. 2020a] through addressing 1) underlying principle that self-regulates the time evolution of whole-genome expression, 2) identification of a peculiar genome region guiding the super-critical genome and determining cell fate change and 3) a unified genomic mechanism to grasp how and when cell-fate change occurs.

## 1.2 Classical Self-Organized Criticality Models

### 1.2.1 c-SOC Model

In physics, the classical self-organized criticality (c-SOC) is a property of dynamical systems that have a critical point as an attractor. This is the natural condition of biological systems at all the organization layers from protein molecules to ecosystems: the equilibrium condition is not a fixed point but a continuous oscillation around a typical configuration (being it a native protein structure or a profile of species relative abundance in an ecosystem) continuously challenged by environmental stochastic variations. The system adapts to these fluctuations, as aptly envisaged in the time-honored physiological concept of homeostasis [Noble 2008] continuously adjusting its configuration, this continuous adjustment is both the basis of the long-term stability of the system and, when exceeding a given threshold, the formal cause of state transitions. This is why c-SOC governed systems reside in the same time in a critical (prone to change) attractor (i.e., stable) state [Huang at al. 2009].

c-SOC governed systems display the spatial and/or temporal scale-invariance characteristic of the critical point of a phase transition. The concept was put forward by Bak, Tang and Wiesenfeld [Bak, 1987,1988], and is considered one of the mechanisms by which complexity arises in nature. c-SOC arises in slowly driven non-equilibrium systems with a large number of degrees of freedom and strongly nonlinear dynamics, where those systems are not drastically perturbed by an external force with continuous external stimuli. This condition can be equated to the ‘normal existence’ of biological systems like tissues or cells in culture exposed to a continuously varying (but dynamically stable) microenvironment.

The paradigmatic c-SOC-based system is the sandpile: think of pouring sand very slowly (ideally one grain at a time), onto a flat, circular surface (this is exactly what we intend for ‘slowly-driven’). At first, the grains stay close to the position where they land, very soon they start to accumulate, creating a pile that has a gentle slope. Going on with the experiment, when the slope becomes too steep, somewhere on the pile, the grains slide down, causing a small avalanche. As we add more sand, the slope of the pile further steepens, and the average size of avalanches increases. The size of avalanches follows a 1/*f* scaling: many small avalanches, very few huge avalanches. The pile stops growing when the amount of sand added balances the amount of sand falling off the edge of the circular surface. At that point, the system reaches the critical state. This state is (dynamically) stable like any proper attractor: the continuous avalanches are counter-balanced by the added sand and the height and shape of the pile remains the same. Nevertheless, occasionally (right part of the 1/*f* spectrum, correspondent to big and consequently very rare events), an added grain can cause a large catastrophic avalanche by a sort of chain reaction involving progressive smaller avalanches falling down until the base and, thus, flattening the entire sandpile (long range correlation, typical of transitional states). The chain reaction can be imagined as a “branching” process, potentially invading large part of the pile (the size of the avalanche can be easily estimated in terms of number of grains involved); in short, each grain falls down until a position of rest, during the slide the grain hits other grains causing small, and then large, avalanches.

It was experimentally demonstrated [Bak and Chen 1991] that, even huge avalanches (posited that we continue to add a grain at a time) keep invariant the slope of the sandpile, given that the probability that a single grain stop is balanced by the probability of a new avalanche. It is worth noting that even huge avalanches invading the entire sandpile happen by a local mechanism (domino effect): each grain only interacts with its neighbor, this is exactly what we observed by RQA as applied to gene disposition along the chromosomes [Zimatore et al. 2021]. If the slope of the sandpile is lower than the critical slope (sub-critical state), the pile will grow until it reaches the critical threshold; if the slope is too steep (super-critical state), the size of the avalanches will be greater than in the critical state so lowering the slope of the pile down to the critical threshold. Thus, the “critical state” attracts both sub-critical and super-critical states. The distribution of the size of avalanches follows a power spectrum distribution: very few large avalanches involving the whole pile, very frequent smaller ones involving only parts of it. In any case, no external observer can predict how, where (in which part of the sandpile) and when a catastrophic avalanche will take place even because the (rare) catastrophic avalanches are ‘caused’ by added grains as small as those provoking very small events. Such “catastrophic” events depend on the past-history of the sandpile and not by the strength of the applied stimulus. This makes c-SOC completely different by other transitions crucially depending on the driving force (control parameter) and question the usual linear relation between the size of the stimulus and its consequences.

Summarizing, the main properties showed by a system governed by c-SOC are:

a. Spatial self-similarity (no characteristic length, small and huge avalanches are both typical of the nature of the ‘stable criticality’ of the system, i.e., they pertain to the same statistical distribution).
b. Temporal persistence (memory effects, the “catastrophic” event depends on its history and not on the applied stimulus).
c. Long-term divergent, correlations (small avalanches together can cause one “catastrophic” event by finding their way across by domino effect).

The domino effect characterizing c-SOC makes it extremely intriguing for explaining the onset of a large avalanche, able to give rise to a huge restructuring of the system at hand. The supercritical state (exactly like in domino), is attained by increasing the density of contacts between the elements (domino tablets), so that a small stimulus can trigger a long and branched chain reaction. Nevertheless, we cannot actively drive such a phenomenon but only setting the conditions fostering the appearance of such chain reaction, whose shape and size will depend only by the structure and history of the system at hand. However, due to this nature, it has been pointed out that fine-tuning of a driving parameter for self-organization in c-SOC generates a controversy regarding the real meaning of self-organization, where there is no need for fine-tuning of an external driving parameter to maintain critical dynamics in SOC control (see more in [Halley at al.2009)]). Furthermore, Sanchez and Newman in plasma physics addressed that c-SOC cannot be considered as a classical physical theory giving rise to exact quantitative predictions [Sanchez and Newman 2015].

### 1.2.2 Extended concept of c-SOC in the cell-fate decision: Rapid SOC model

Halley et al. [Halley et al. 2009] extended the concept of c-SOC so to model the cell-fate decision (critical-like self-organization or rapid SOC) through the extension of minimalistic models of cellular behavior. A basic principle for the cell-fate decision-making is that gene regulatory networks adopt an exploratory process, where diverse cell-fate options stem by pruning of various transcriptional programs. Then a cell-fate gene module is selectively amplified as the network system approaches a critical state; these review articles also present a useful survey of studies on self-organization in biological systems.

Self-organizing critical dynamics of this type are possible at the edge between order and chaos and often go together with the generation of exotic patterns. Self-organization is considered to occur at the edge of chaos [Langton 1990; Kauffman 1993] through a phase transition from a subcritical domain to a supercritical domain: the stochastic perturbations initially propagate locally (i.e., in a sub-critical state). Thus, due to the particularity of the disturbance, the perturbation can spread over the entire system in an autocatalytic manner (into a super-critical state) and thus, global collective behavior for self-organization develops as the system approaches its critical point.

## 1.3 Self-Organized Critical Control (SOC control) of Genome Expression Regulation

Through our previous studies [Tsuchiya et al. 2014, 2015, 2016, 2017, 2020a; and Giuliani et al., 2018], we identified a peculiar set of genes (i.e., critical point, CP) that guides the super-critical subset. CP is where coordinated global perturbation starts. In scientific literature, we were able to find distinct cell-fate change processes, ranging from single cells to population dynamics following the above-sketched model. Namely: HRG-stimulated MCF-7 cells [Nagashima et al. 2007; Nakakuki et al. 2010], atRA- and DMSO-stimulated HL-60 cells [Huang et al. 2005], Th17 cell differentiation from Th0 cell [Ciofani et al. 2012], mouse and human early embryo development ([Deng et al. 2014] and [Yan et al. 2013]).

The above cases are examples of a general mechanism and have been selected based on the availability of a sufficient number of experimental time points.

Figure 1 reports a graphic summary of the proposed mechanism of cell-fate change. The analysis of real biological data forced us to modify classical c-SOC introducing six relevant changes:

1. *Hot Spots*: Each cell-type has a specific and largely invariant (at the entire genome scale) gene expression profile. The continuous adaptation to microenvironment fluctuations happens by the agency of a set of genes with high variance in time responsible for the scattering across the identity line of gene expression at different times. This situation is mirrored by the presence of a main first principal component (PC1 accounting for around 90% of total variance) correspondent to the attractor in the gene expression space (GA) and by two minor (PC2, PC3) components, orthogonal to PC1 accounting for fluctuations around equilibrium position. We call ‘hot spots’ the genes with higher scores on these minor fluctuation modes [Zimatore et al 2021]. When the entity of variance explained by minor components exceeds a critical threshold, the fluctuation is transmitted to the core of the system and the transition can start. Hot spots play the role of continuous avalanches of sandpile that occasionally provoke the transition when their motion invades the entire system by a domino effect activating the CP.
2. *Critical Point (CP)*: Our findings of SOC does not correspond to a phase transition in the overall expression from one critical state to another (i.e., sub-critical to super-critical genome state transition), describing the genome approaching to a critical point (CP) as developed in c-SOC. Instead, whole-genome expression is dynamically self-organized through the emergence of a CP, with the co-existence of three distinct response domains (local critical states). The temporal variance of expression (normalized root mean square fluctuation (*nrmsf*); see Methods) acts as an order parameter for the self-organization of whole-genome expression [Tsuchiya et al. 2015, 2016]. The properties of the CP such as sandpile critical and scaling-divergent behaviors were described in details in our previous paper (refer to Fig. 4 in [Tsuchiya et al. 2016]). Note: our approach on genome expression is based on physical mean-field theory (https://en.wikipedia.org/wiki/Mean-field_theory). Our studies [Censi et al. 2011; Tsuchiya et al. 2015] have demonstrated that the law of large numbers tends to hold when more than 50 genes are randomly selected: its ensemble average converges to its center of mass. This feature (elimination of expression noise in individual genes) is crucial to obtain the essential behavior on the dynamics of whole-genome; i.e., the dynamics on the ‘center of mass’ deduced from the ensemble gene average (gene mean-field) provides reliable insights into the behavior of the genome system. Thus, we revealed self-organized critical control (SOC) of whole expression to study dynamics in relation to the higher-order fluctuations of gene expression.
3. *State Change in the CP*: The ON-OFF switch of the CP state occurs through change in critical transition on its singular-bimodal behaviors (Figure 1C). The CP represents a specific set of critical genes, which has an activated (ON) or repressed (OFF) state. It is to be noted that this ON-OFF state does not indicate a simple binary state switch, but admits different state levels. Furthermore, this critical-genes (CP) induces a change in genome attractor (GA, center of mass of whole expression). The synchronization of expression dynamics between the CP and GA occurs at the critical transition in HRG response, whereas in EGF response, no such synchronization occurs (Figure 1B). This synchronization suggests how the change in the CP induces change in the GA and thus, induces global expression avalanche (Figure 1D).
4. *Genome-Engine*: Open-thermodynamic dynamic picture of between-state flux provides a potential universal mechanism of self-organization interpreted in terms of a “genome engine” (Figure 1E). An autonomous critical-control genomic system is developed through the highly coherent behavior of low-variance genes (local sub-critical state), which in turn, generates a dominant cyclic expression flux with high-variance genes (local super-critical state). This became evident after gene expression was sorted and grouped according to the order parameter (*nrmsf*). When whole expression is sorted and grouped according to the order parameter, gene mean-field approach reveals the emergence of its self-organization with critical transitional behaviors such as sandpile type and scaling divergent ones (e.g., Fig. 4 in [Tsuchiya et al. 2016]). On the contrary, randomly shuffled gene expression exhibits no evidence of cooperative behavior (no emergence of coordinated activity).
5. *Switching of Genome-Engine*: Average value of external expression flux into the genome expression system is negligible if compared to internal expression flux between local critical states. However, in terms of fluctuation from the mean genome-engine, external flux from the cell environment is not small at all, especially being large around critical transitions for cell-fate change (see section 1.4), suggesting that symmetry breaking of the cell-fate change reaches peak around the critical transition. Thus, inflow and outflow of expression flux to the genome system is balanced. Due to this balance, the law of large numbers holds: as the number of elements increases, average value of randomly selected expression converges to that of whole expression, i.e., this is the biophysical reason for the existence of the GA. Interestingly, coherent switching of the genome-engine occurs through the cell-fate guiding critical transition (Figure 1F; see more in section 1.4), where this genome-engine switching is driving force for the cell-fate change.
6. *CP acting as the Center of Cell-fate Change*: Cell-fate change occurs through erasure of the initial-state sandpile critical behavior (criticality), where the corresponding initial-state global gene expression regulation such as epigenetic modifications and chromatin remodeling is eliminated. Thus, the CP acts as the center of cell-fate change, where its state-change guides the genome from a sub-critical to a super-critical genome state.

**Figure 1:**
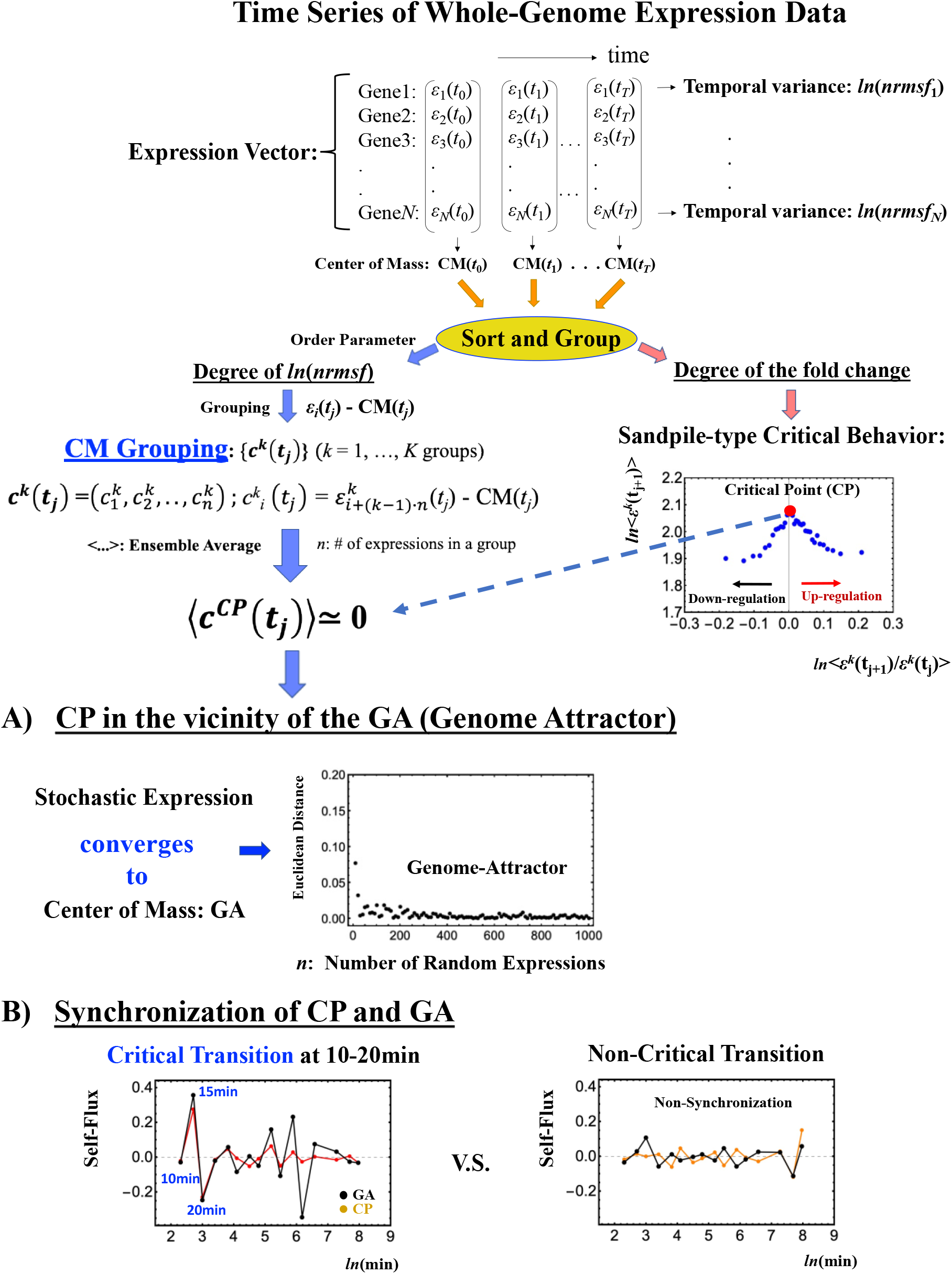
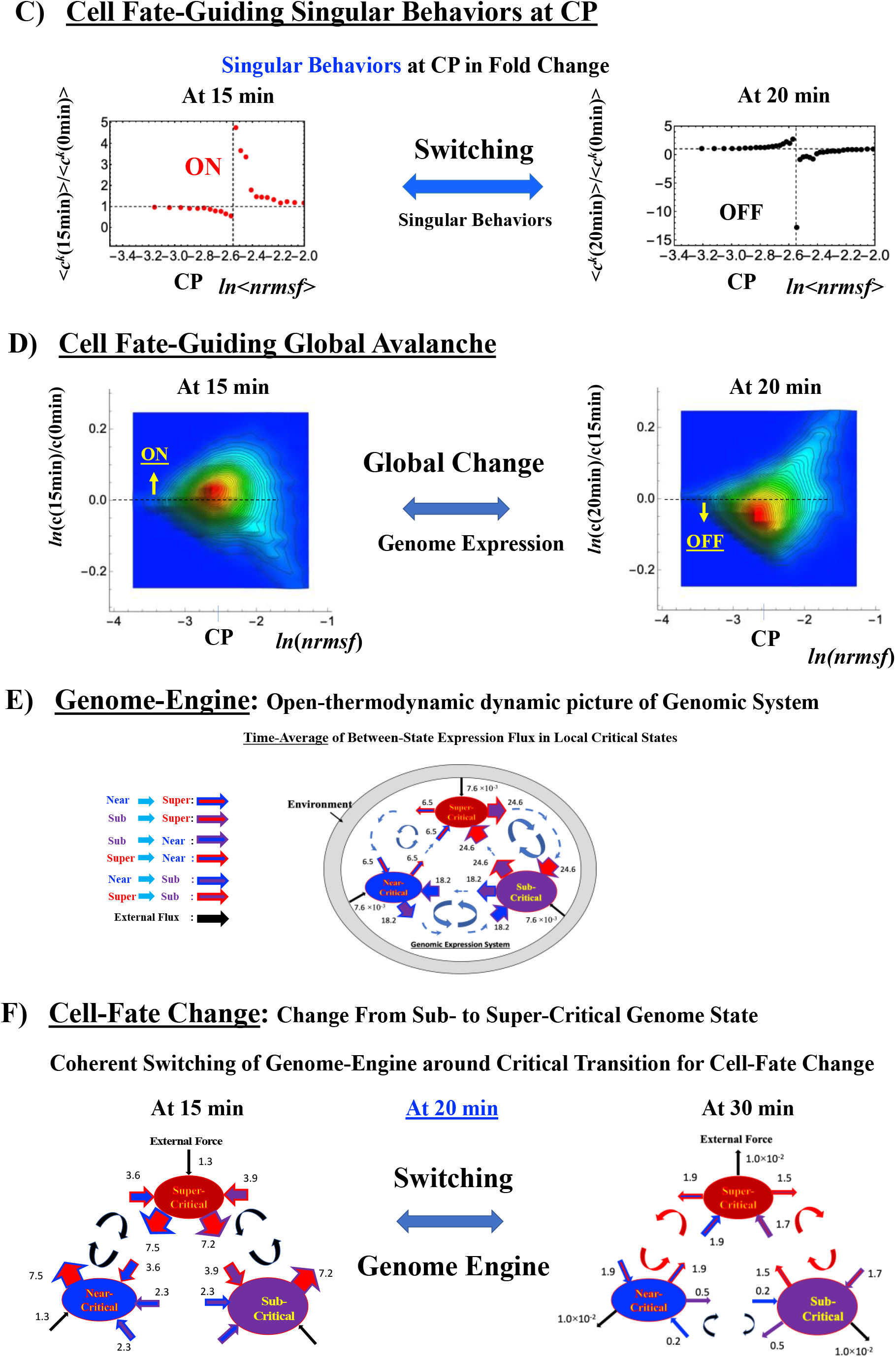
Updated unified genomic mechanism for cell-fate change: Self-Organized Critical (SOC) control of time-series whole expression for cell-fate change. A unified genomic mechanism is described by MCF-7 cell response as an example (refer to Fig. 4 in [Tsuchiya el al. 2020a]). The GA corresponds to the center of mass (CM) of whole gene expression and acts as genome-attractor. **A)** The CP, where the CP is the summit of sandpile critical behavior, exists in the vicinity of the GA (see Fig. 2 in [Tsuchiya el al. 2020a]). **B)** Synchronization between the CP and GA (critical transition at 10-20min; left panel) occurs for HRG-stimulated MCF-7 cells (cell-differentiation), whereas for EGF-stimulated MCF-7 cells, such synchronization does not occur (right panel). **C)** Switching of critical transition at the CP with bimodal expression distribution reminiscent of swelled coil (ON) and compact globule (OFF) DNA transition corresponds to the inversion of the intra-chain phase segregation of coil and globule states. **D)** When the switching at the CP, global avalanche of gene expression happens: at OFF of the CP, density profile of whole expression (probability density function: PDF) shifts towards down-regulation, whereas for ON of the CP, PDF shifts towards up-regulation. **E)** Updated genome engine mechanism (see section 1.4 and Methods) revealed through average between-state expression flux. CM of a critical state acts as a local attractor (see Fig. 10 in [Tsuchiya et al. 2016]). SOC of whole expression emerges as a self-organized heteroclinic (more than one attractor) critical control to guide collective (coherent) behaviors emerged in the critical states. **F)** Temporal fluctuation on the gnome-engine reveals that around cell-fate change, coherent switching on the genome-engine occurs (see section 1.4).

Regarding material basis of SOC control, Krigerts et al. [Krigerts et al. 2021] revealed that the scale-free pulsing of the pericentromeric domains (PADs) likely erasing their differential gene-silencing position effect at 15 min after HRG treatment coincides in time with high wave of the gene expression dynamics started with the CP gene group of early stress response possessing bi-valent chromatin structure enabling bi-phasic (up-and-down activity in the time course). This correspondence in time between PADs bursting and gene expression dynamics is the most cogent proof of the material basis of the SOC model. The establishment of a link between gene expression dynamics and experimental observation of structural changes in chromatin organization is of utmost importance, because it gives a material basis to the proposed model and identifies chromatin folding/unfolding dynamics as the main player in the large-scale gene expression regulation. The specificity of the large-scale regulation is not guided by the ‘intelligent agents’ but by the symmetry breaking of the nuclear space induced by the topology of chromatin networks [Krigerts et al. 2021; Erenpreisa et al. 2021], so fulfilling the conditions for the generation of order from chaos predicted by non-linear thermodynamics [Erenpreisa and Giuliani 2020].

## 1.4 Genome-Engine: Open-thermodynamic View of Genome Expression System

In this section, here we update genome-engine as we understand it [Tsuchiya et al. 2020a]. Four crucial points are listed as below:

1. The CP is different from the GA (more in the next section), where we described that the CP acts as the GA. The CP is located in the vicinity of the GA (Figure 1A), where the CP and GA form a phase segregation (bimodal domains). Interestingly, any part of the CP in chromosome expresses similar dynamic behavior to the whole expression of the CP (see section 1.6), i.e., exhibits scaling behavior (renormalization).
2. The state-change in the CP induces critical transition to guide global expression avalanche (see details in [Tsuchiya et al. 2020a]). The CP and GA are synchronized around the critical transition (Figure 1B: see more in section 1.6), which derives the change in the GA and thus, global perturbation occurs. On the other hand, in EGF-stimulated MCF-7 cells (only cell-proliferation: no cell-fate change), critical transition does not occur due to the fact that CP and GA are not synchronized in time.
3. Critical behaviors at the CP exhibits bimodal behaviors of activation (unfolded swollen chromatin) and suppression (folded compact chromatin), where at critical transition switching of singular behaviors (that is a small change causes large effect) occurs (Figure 1C). Physically, the appearance of the critical behavior can be described in terms of the depression of the free energy barrier between the active and silent genome compartments. In statistical physics, it is well-established that criticality appears accompanied by the depression of the free energy barrier in the bimodal free energy profile for the system of first-order phase-transition. In relation to this, it has been confirmed that genome-sized DNA above the scale of several tens kp exhibits the characteristics to undergo folding transition from elongated state into compact state in an all-or-none manner [Yoshikawa et al. 1997, 2002b, 2002c; Tsumoto et al. 2003; Yamada et al. 2005; Krotova et al. 2010; Tongu et al. 2016]. Near criticality, fluctuation of the state of bimodal phases or phase separation becomes significant, and scaling-like behavior appears. From experimental and theoretical studies [Iwataki et al. 2000; Takagi et al. 2001; Iwaki and Yoshikawa 2003; Zinchenko et al. 2003; Nakai et al. 2005; Takenaka et al. 2008; Nagahara and Yoshikawa. 2010; Yoshikawa et al. 2011], it is getting clearer that, for genome-sized DNAs, critical behavior emerges through the competitive effect among conformational entropy of DNA, histone modification, nucleosome stability and translational entropy of environmental ionic species including mono and multivalent cations, ATP, RNA. Here, it is noted that the critical behavior as well as bimodal free-energy profile is the specific characteristics for genome-sized DNA and that such characteristics disappear for shorter DNA less the scale of several kbp. In HRG-stimulated MCF-7 cells, possible biological mechanism of depressing free energy barrier might stem from spatial interaction of two chromatin types - silencing pericentric (hetero-) chromatin domains (PADs) and transcribing euchromatin. In control, the large PAD clusters labelled by H3K9me3-marked repressive heterochromatin are surrounded by a ‘collar” of intermingled the repressive and active H3K9me3+H3K4me3 marks. However, at 15 min after HRG, this barrier is released and the active transcribing H3K4me3-marked euchromatin becomes repulsed from split smaller PADs (see Supplementary S1 in [Krigerts et al. 2021]). Due to lifting of the entropy barrier, expression dynamics of the CP and GA can synchronize to guide the global expression avalanche for cell-fate change. PADs change was accompanied by simultaneous and sustainable 1.5-fold (p< 0.01) increase of DNA unfolding (determined by the Acridine orange DNA structural test) and 2-fold (p< 0.01) increase of homogeneity of the euchromatin distribution in cell nuclei [Krigerts et al. 2021].
4. Expression flux (second order time-difference of gene expression) at a discrete time point can be exactly described as expanded in internal-interaction and external-force terms, where internal interaction terms reveal how expression flux form a cycle-state flux, while external flux (force) term represents how much energy is coming or going out to the genome system through the cell nucleus environment (see Methods).

With these updates, the key point of genome engine mechanism is that dynamics of distinct coherent behavior emerges from stochastic expression in local critical states, i.e., coherent-stochastic behavior (CSB). Center of mass (CM) of any randomly selected gene expression from a critical state converges to the dynamics of CM of the critical state, which means that the CM of a critical-state acts as the critical-state attractor. Genome-engine represents overall picture of time-average between-state flux (Figure 2) among three critical-state attractors (heteroclinic genomic system), which is a stable manifold of the thermodynamically open system (see more in Methods).

**Figure 2:**
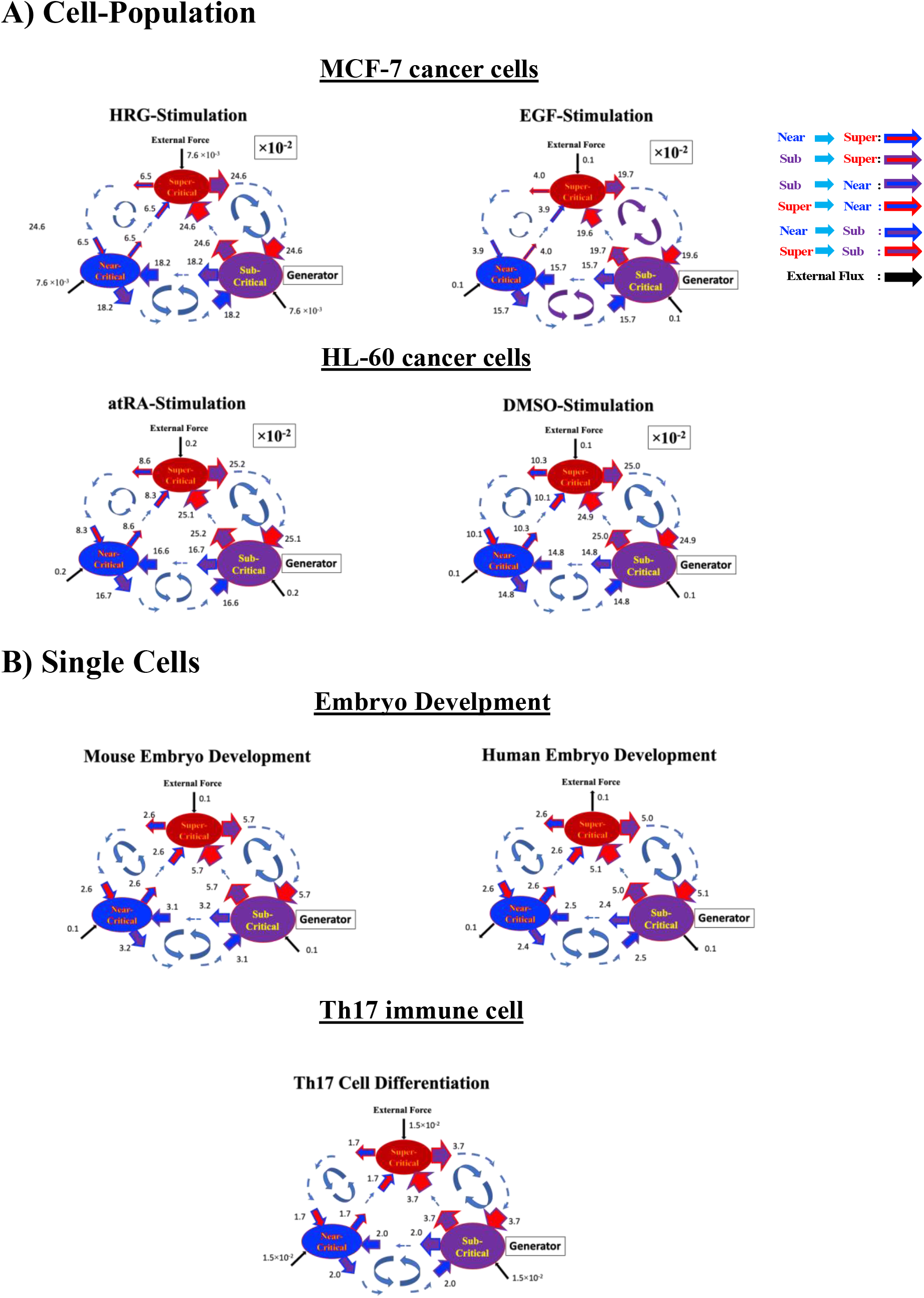
Updated genome-engine mechanism. Sub-super cyclic flux forms a dominant flow, where the sub-critical state acts as generator of the genome engine with sending large fluxes from the sub-critical to other critical states. The CP acts as an “ignition switch”, where change in the CP spreads through these cyclic fluxes. The cyclic flux between sub-critical state and another critical state derives the anti-phase dynamics of two critical states. Cell-Population: **A)** HRG-stimulated (left top) and EGF-stimulated (right top) MCF-7 cancer cells; atRA-stimulated (left bottom) and DMSO-stimulated (right bottom) HL-60 cancer cells. Single-Cell**: B)** mouse- (left top) and human-embryo (right top) development; Th17 immune cell development (center bottom).: Numerical values represent average between-state expression flux, whereas in the cell population, the values are based on 10^-2^ scale.

We summarize the systemic view about how and when cell-fate change occurs, as follows:

### 1) How cell-fate change occurs

Coherent switching of genome-engine in terms of interaction fluxes (Figure 3) between critical states occurs before and after the cell-fate change. Thermodynamically, the genomic system passes over a non-equilibrium fixed point through a large change in external force. This coherent switching shows that a global perturbation around the cell-fate change induces enhancement or suppression on the genome-engine, where there is a dominant cyclic flow between the super- and sub-critical states (see Tables 1, 2 in [Tsuchiya at al 2020a]). In HL-60 cells (cell population) treated with DMSO, the genome-engine is enhanced before the cell-fate change and suppressed (enhancement-suppression) thereafter, where genome-engine for DMSO stimulation receives the ‘biological feedback’ from cell nucleus environment, whereas reverse process occurs for atRA stimulation (Figures 3B, C). On the contrary, a reverse process of suppression-enhancement takes place in the MCF-7 cancer cells undergoing the commitment to differentiation (Figure 3). In these cases, perhaps the time interval between commitment and determination, as well as the time-dependent volume of the involved in the process cell population may play role [Krigerts et al. 2021]. In the single-cell cases (embryo development and Th17 immune cell), a suppression-enhancement on the genome engine occurs (Figure 4). The varying sequences of perturbation on the genome-engine may stem from different stages of the suppressive pressure on cell-differentiation against cell-proliferation.

**Figure 3:**
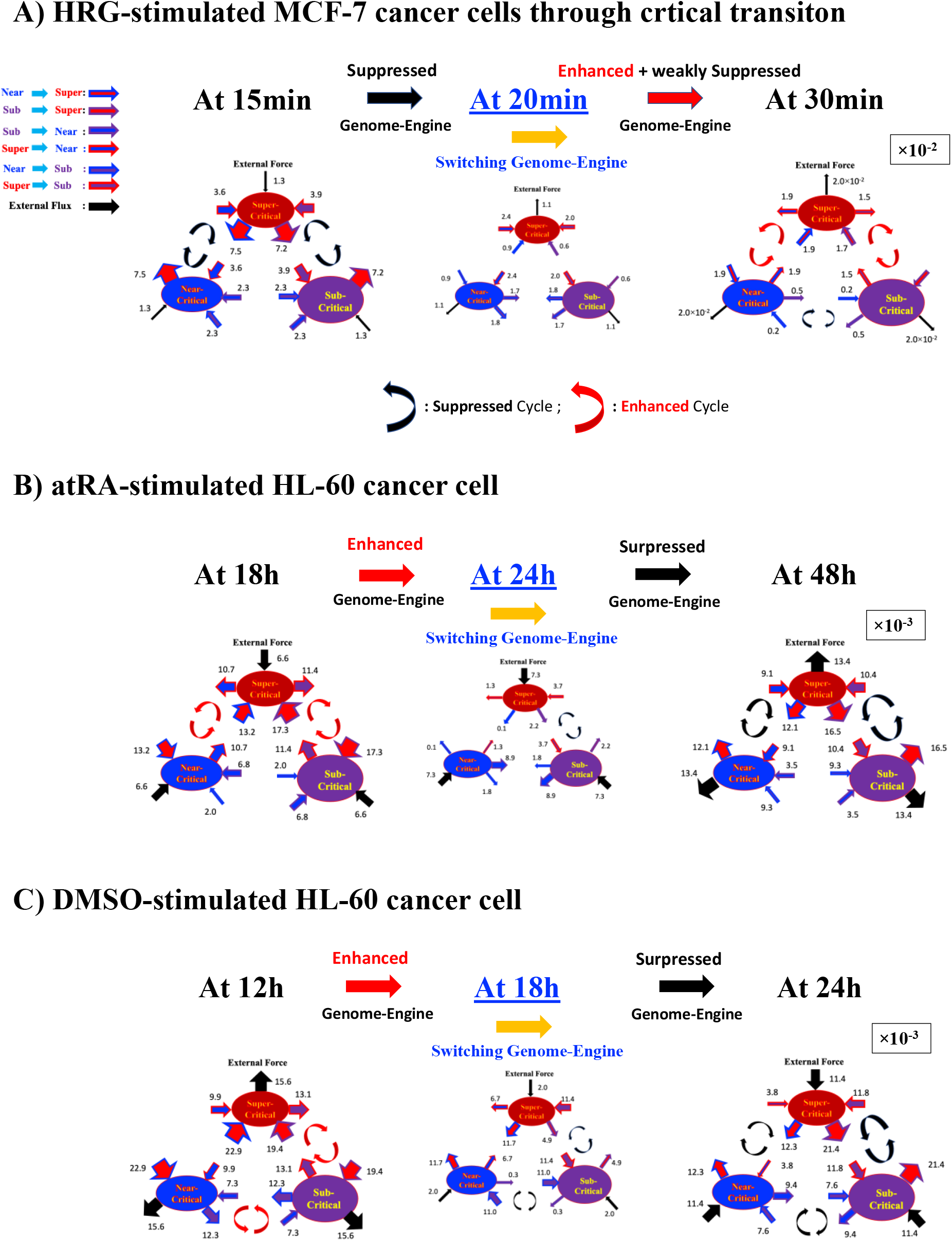
Cell population - coherent switching on the genome engine through cell-fate change. Cell fate mechanism through perturbation on the genome-engine: **A)** MCF-7 cells; **B)** HL-60 cells. Before the cell-fate change, the onset of the switch to coherent dynamics occurs on the dominant cyclic flow: switching from suppression to enhancement for MCF-7 cells, whereas switching from enhancement to suppression for HL-60 cells. The timing of coherent switching on the genome-engine comes along with that of cell-fate change (see Tables 3,4 in [Tsuchiya et al. 2020a]). Numerical values based on 10^-2^ for MCF-7 cells and 10^-3^ scale for HL-60 cells represent interaction flux. Refer to Methods for further detail experimental time points.

**Figure 4:**
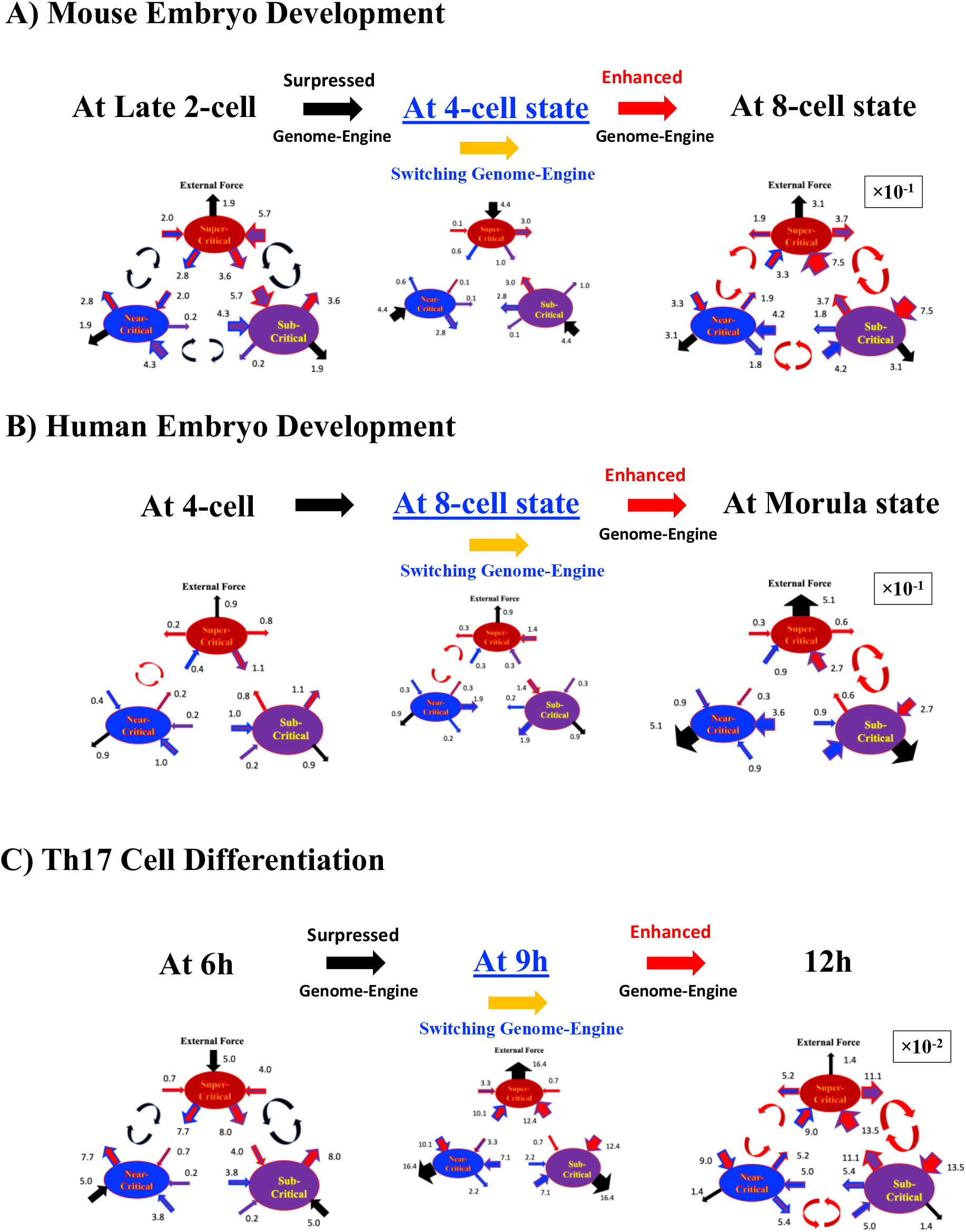
Single cell - coherent switching on the genome engine through cell-fate change: **A)** Mouse embryo development**; B)** Human embryo development; **C)** Th17 immune cell differentiation. Reprogramming on embryo development and cell differentiation on Th17 immune cell occurs through the suppression-enhancement on the dominant cycle state flux between super- and sub-critical states, which is opposite to the cell differentiation in HL-60 cells. The timing of coherent switching on the genome-engine comes along with that of cell-fate change (see Tables 3,4 in [Tsuchiya et al. 2020a]). Numerical values based on 10^-1^ for embryo cell and 10^-2^ scale for Th17 cell represent between-state expression flux.

### 2) When cell-fate change occurs

Cell-fate change corresponds to the erasure of initial-state sandpile criticality (i.e., erasure of the initial-state CP memory). That is to say that, before entering a new stable state, the system must eliminate the ‘adaptive memory’ of the previous condition. This is particularly evident in embryo development, in which the new entity (embryo) must first erase the regulatory state of the egg (see Fig.4 in [Giuliani et al. 2018]). The coherent switching to another regulation state occurs on the genome-engine (global perturbation), where the timing coincides with that of the cell-fate change (see Tables 3,4 in [Tsuchiya et al. 2020a]). This suggests that the cell-fate guiding change induced by change in the CP state corresponds to the erasure of initial-state sandpile criticality. In HRG-stimulated MCF-7 cancer cells, however, time lag reveals between global perturbation committing for differentiation (at 15-20min) and cell-fate determination (at 3h).

Therefore, it will be essential for cell-fate determination to understand how activation-deactivation mechanism for the CP uncovers through a coordinate chromatin structure change [Zimatore et al. 2021] partially described by [Saeki et al. 2009; Krigerts et al. 2021]. Furthermore, our SOC hypothesis is expected to predict when and how a cell-fate change occurs in different biological situations (e.g., iPS cells and genomic reason for not reprogramming). The proposed universality of the SOC stems from the existence of very basic physical constraints (e.g., chromatin dimension and general organization, phase transition phenomenology) independent of biological specificities.

## 1.5 Self-Organization of Whole Gene Expression through Coordinated Chromatin Structural Transition

It is worth noting that the above processes are purely local, i.e., the expression activation-suppression flow is transmitted across neighboring tracts of the genome, as demonstrated in [Zimatore et al. 2021], consistently with both the domino effect at the basis of SOC hypothesis and the observed changes in chromatin organization [Krigerts et al. 2021]. Figure 5 reports the co-expression level (*y*-axis) of gene pairs as a function of their spatial distance along the chromosome: the exponential decay of co-expression is a clear proof of the local character of regulation [Zimatore et al. 2021]. The discrimination between effective and non-effective stimuli is immediately grasped by a principal component analysis (PCA) of the matrix having as statistical units the genes and as variables the different times. The presence of a strong, cell-type specific gene expression profile, generates a first principal component (PC1) explaining the by far major part of data set variance. This variance is due to the presence of large difference between genes average expression level, in statistical terms it is a ‘between genes’ variance. The second and third components, explaining a minor part of data set variability [Zimatore et al. 2021], account for ‘within genes’ variance, i.e., for the fluctuation around the ‘equilibrium position’ of each gene expression. Figure 6 clarifies this point showing how, at the global genomic scale, the gene expression profile of two cell populations correspondent to different time points are each other near unity correlated (thus giving rise to a first component accounting for the by far major proportion of between gene expression variance), while the within gene variance (temporal variability) orthogonal tom PC1, explains much smaller part of total variability.

**Figure 5:**
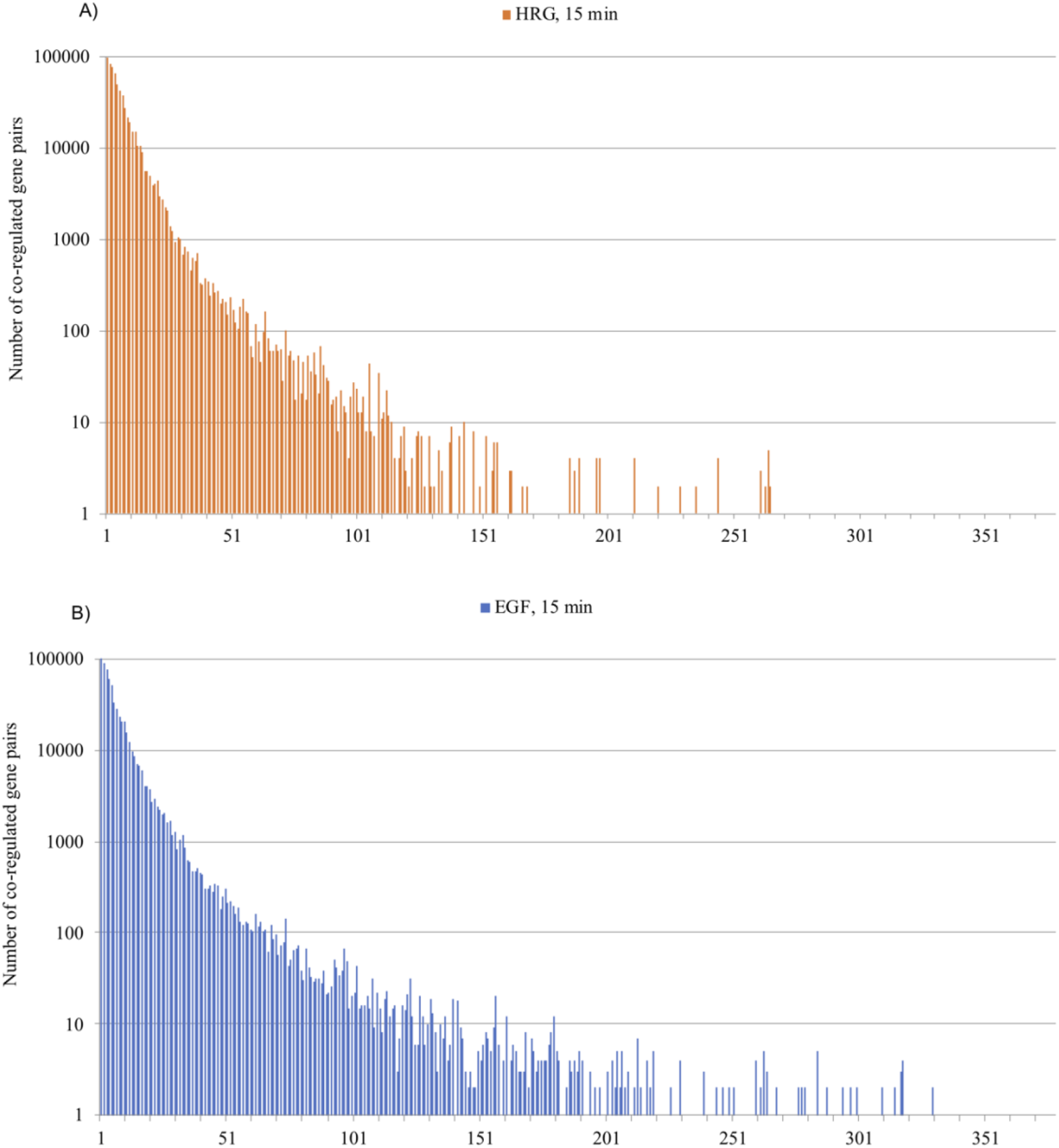
Recurrence spectrum along the chromosome 1 cited from [Zimatore et al. 2021]. The figure refers to MCF-7 case and it is evident how the locality stems from an intrinsic structural constraint (chromatin structure), being present in both effective (HRG stimulation, top panel) and non-effective (EGF stimulation, bottom panel). The graphs are relative to HRG (**A**: top) and EGF (**B**: bottom) conditions as for time 15 min and chromosome 1. The y-axis corresponds to the number of recurrences (number of co-regulated gene pairs). The x-axis corresponds to the spacing of recurrent couples along the chromosome. The spacing is expressed in terms of number of genes between the recurrent pairs. The exponential decay of co-regulation entity with increasing between genes spacing is identical for different times of observation, different chromosomes and for the two EGF and HRG conditions. The local character of fine co-regulation around cell-kind specific value is the same for all the experiments.

**Figure 6:**
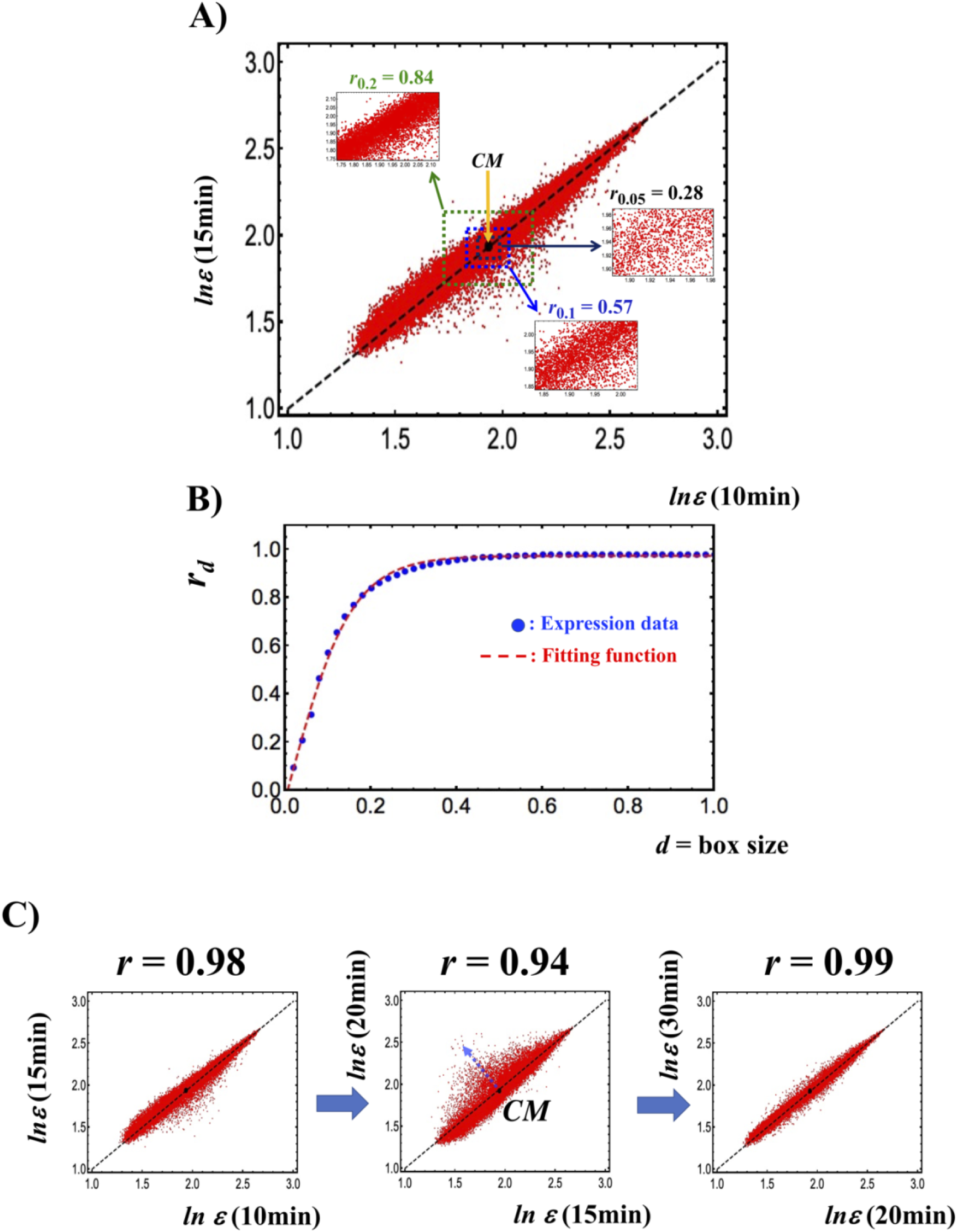
Transition from a stochastic to a genome-wide attractor profile cited from [Tsuchiya et al. 2016; Zimatore et al. 2021]. **A)** Pearson’s correlation between two independent samples (10min vs. 15min) of MCF7 (axes of the plot) in terms of expression levels of around 23000 mRNAs (vector points). **B)** Profiles correlation rd versus d (box size), see the tangent hyperbolic function as signature of transition from a stochastic to a genome-wide attractor profile. **C)** The Pearson’s correlation between gene expressions profiles at different time points reveals the occurrence of critical transition at 15-20 min around the CM (back solid dot) correspondent to a decrease in Pearson r and visually to an increase scattering of points from the identity line. A tipping point in terms of the critical transition exists at 15min (see more in Figure 5). This stems from the (coil-globule) critical transition at the CP, where the CP corresponds to the CM when expression is ordered by nrmsf. The clear identity line between gene expression profiles at different times is the image in light of the existence of a major principal component (PC1). The scattering orthogonal to such identity line corresponds to the minor ‘motion’ components (PC2, PC3). This scattering (and consequently the percentage of variance explained by the minor modes) is maximal at transition (central graph of panel C).

The within-gene temporal variability corresponds to the ‘critical attractor’ of SOC driven process: the small fluctuations needed to keep the genome expression at the pace of external unpredictable small perturbations. Minor components are the image in light of physiological homeostasis, in the same time, these motion components are at the basis of the occurrence of catastrophic avalanches in the rare case when their power (amount of variance) exceeds a given threshold. This is exactly what happens in the HRG (efficient stimulus) case and does not happen in the EGF condition: 15 minutes after treatment, the percent of variance accounted by minor (motion) components is 34% in the HRG and 10% in the EGF case, this means that the HRG-induced perturbation (at odds with EGF) is sufficient to ignite the transition [Zimatore et al. 2021].

## 1.6 Synchronization Between Critical Point and Genome Attractor: CP as Organizing Center of Cell-fate Change

As described in Figure 1A, taking CM of genome expression (i.e., the GA) as baseline, the CP exists in the vicinity of the GA (see more in Fig. 2 in [Tsuchiya et al. 2020a]). As shown in whole gene expression analysis in HRG-stimulated MCF-7 cells compared with non-differentiated EGF-stimulated ones, the state change of the CP occurs in the HRG response, whereas in the ERG response, that of the CP does not occur (Fig. 4 in in [Tsuchiya et al. 2020a]; confirmed further by repeated data). Figure 1B is to show that synchronization between the CP and the GA occurs in the HRG response at 10-20min (critical transition), whereas no synchronization occurs in the EGF response. This synchronization in terms of average behavior (mean-field behavior) describes how the change in the GA occurs to induce genome avalanche in the genome.

### 1.6.1 Critical Transition Transmitted to the Genome Through Synchronized Chromosomes with the CP dynamics

Next, we investigate how and where the synchronization transmits in chromosome level through the critical transition. Notably, Figure 7 reveals that any portion (average behavior) of the CP belonging to a chromosome follows almost the same temporal expression response as the dynamics of the whole CP, which describes scaling behavior (renormalization) in the CP genes in chromosome level. Furthermore, in chromosome level, dynamics of a chromosome (average behavior) synchronizes with the portion of the CP genes at the critical transition. Coherent-stochastic behavior (CBS; see [Tsuchiya et al. 2015]) also occurs in a chromosome level (not Figure shown here), where stochastic gene expression within a chromosome converges to its CM of expression (as a local attractor), such that the principle of synchronization in chromosome level also stems from the synchronization with the CP.

**Figure 7:**
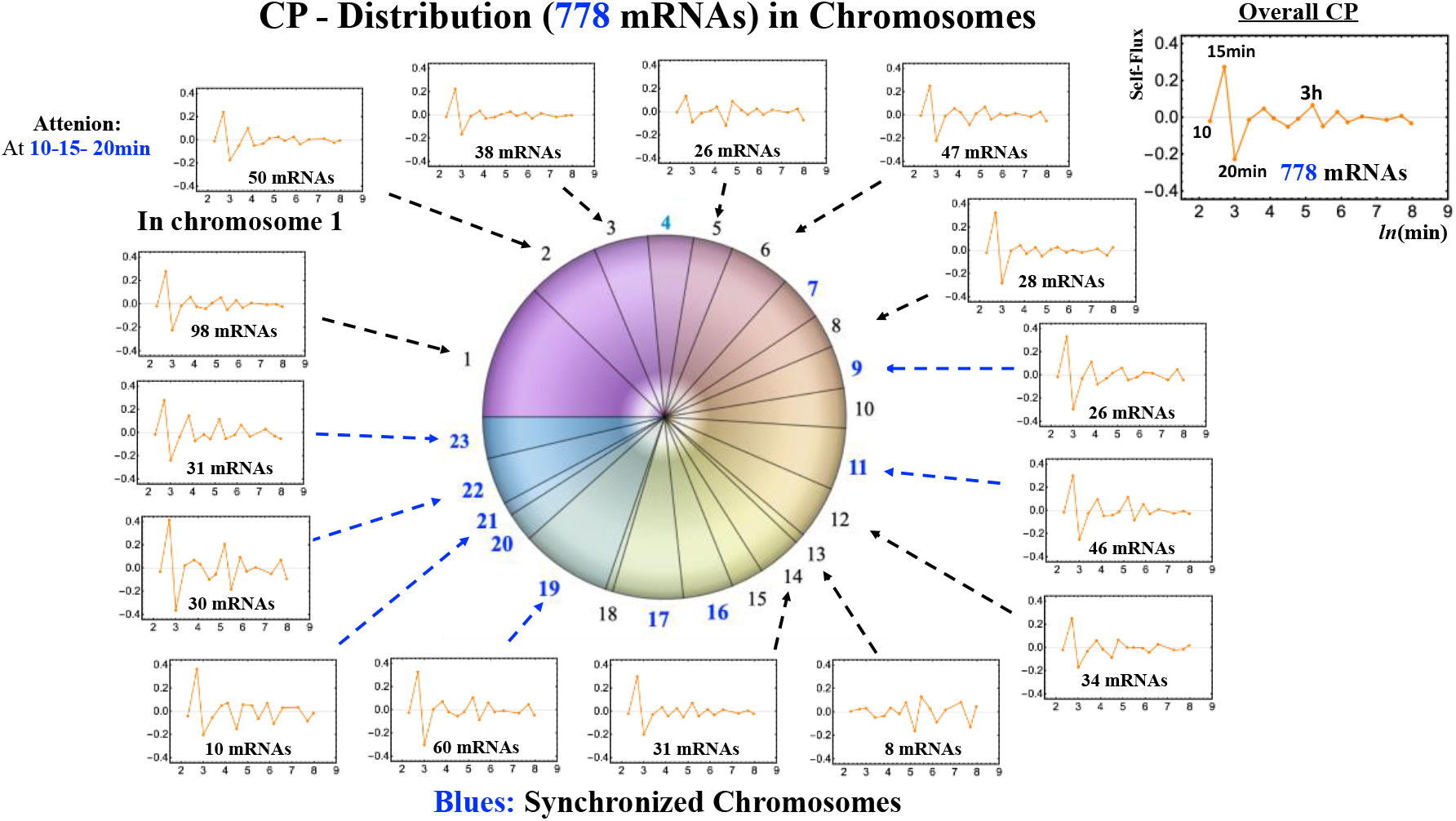
Distribution of the CP genes into chromosomes: Temporal behaviors of the CP genes in chromosomes follow overall flux behavior of the CP (778 mRNAs for -2.60 < *ln*(*nrmsf*) <-2.56), which reveals scaling behavior (renormalization) in the CP genes, suggesting the formation nucleation in the vicinity of the GA.

Interestingly, there exist synchronized and non-synchronized chromosomes with the CP (Figure 8). Clustering of synchronized and non-synchronized chromosomes (Figure 9) shows how and where the critical transition transmits to the genome. The GA is located at the branching of synchronized chromosomes and moreover, self-flux of the GA is amplified to transmit through two paths, one from the GA to chromosome 22, and the other from the GA to chromosome 17 (Figure 10). Here, self-flux is the second order time-difference of gene expression to represent flux of “force”, in which force can be interpreted by “biophysical energy” transferring to a chromosome for a positive sign and released from the chromosome for a negative sign (see more in Methods). These paths reveal spatio-temporal behavior of the critical transition with amplified energy transfer in the cancer chromosome territory (MCF-7 cancer cells).

**Figure 8:**
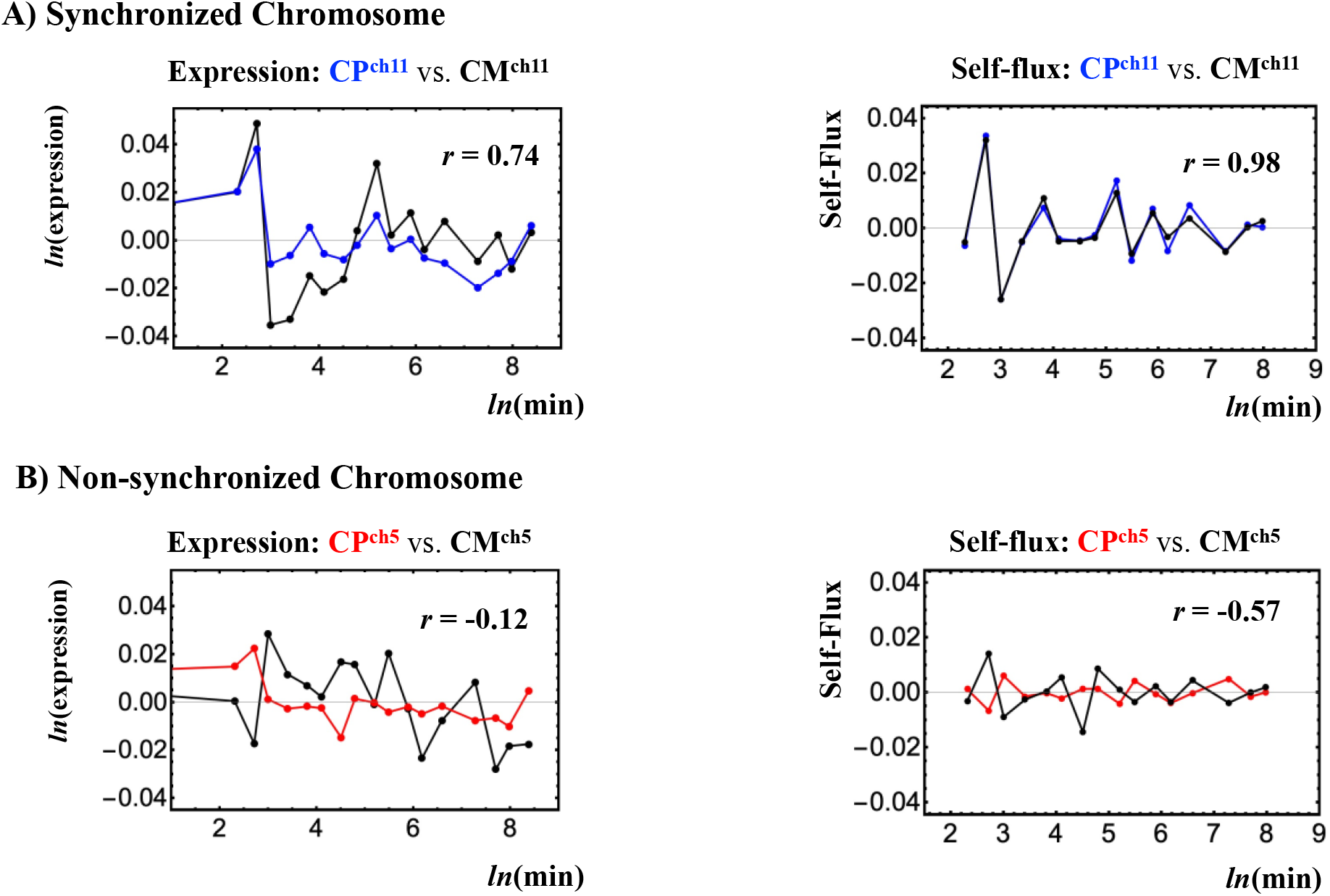
Synchronized chromosomes versus non-synchronized chromosomes in HRG-stimulated MCF-7 cells. **A)** Chromosome 11 (Left panel): ensemble average of expression has a good temporal Pearson correlation (*r*) to the corresponding CP (chromosome-CP), whereas self-flux (second order time difference: right panel: see more in Methods) show a clear synchronization between chromosome and CP. This is a reason to adopt self-flux for synchronization. **B)** Chromosome 5: no synchronization occurs between chromosome and CP. Clusters of synchronized and non-synchronized chromosomes are given in Figure 9. Numerical values of self-flux of chromosome 11 CP (panel A: blue line in right figure) are magnified by three times. The *x*-axis represents natural log of experimental time points in minutes (refer to Methods).

**Figure 9:**
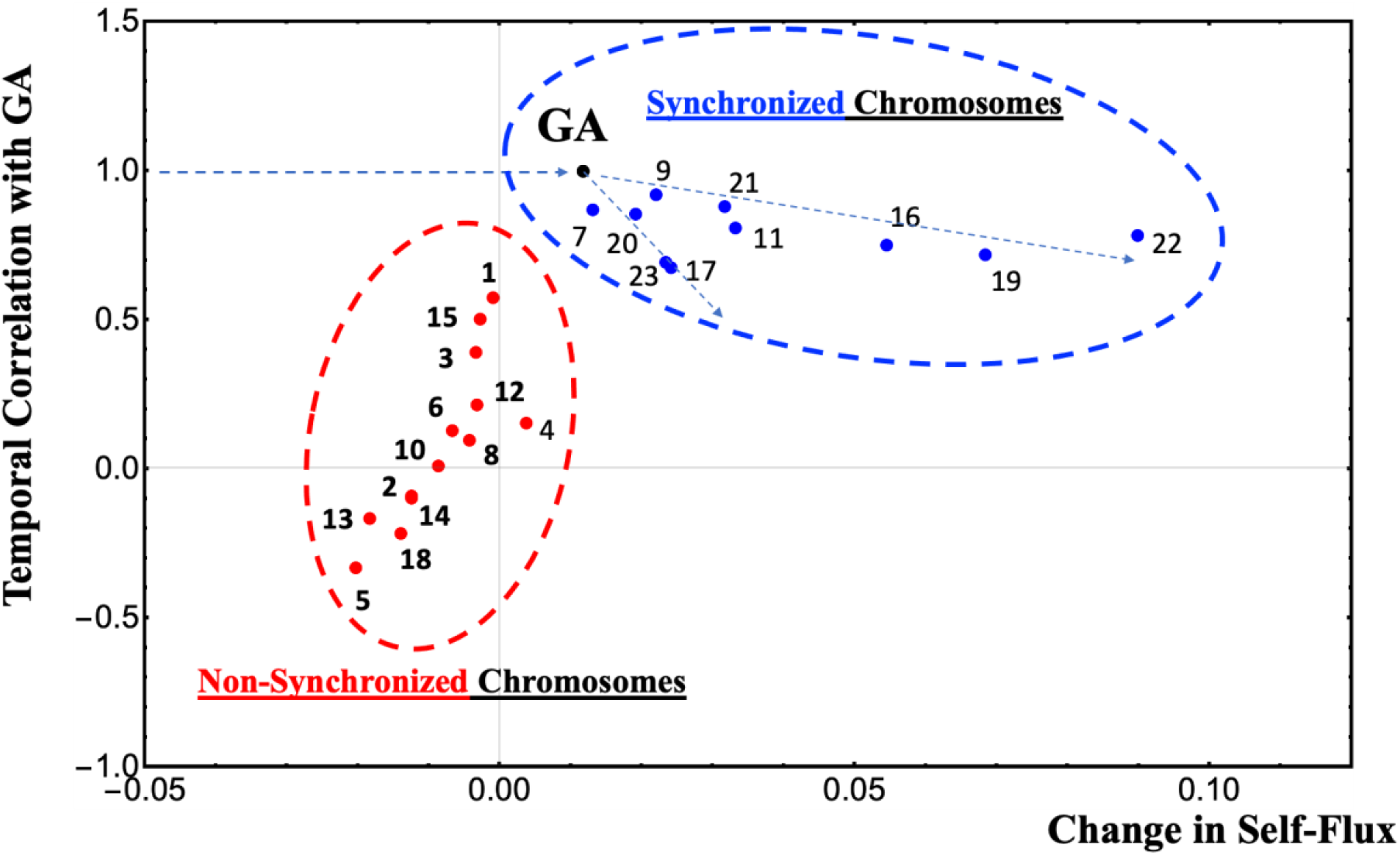
Clusters of synchronized chromosomes and non-synchronized chromosomes. The *x*-axis represents the change in self-flux between 10 to 15 min and 15 to 20 min, and the *y*-axis represents temporal Pearson correlation of CM expression of chromosome with the GA. Plot shows clusters of synchronized chromosomes and non-synchronized chromosomes with the CP. The GA is located at the branching of synchronized chromosomes, which shows how and where the critical transition at 10-20 min transmits through the chromosome territory of MCF-7 cancer cell genome (see Figure 10).

**Figure 10:**
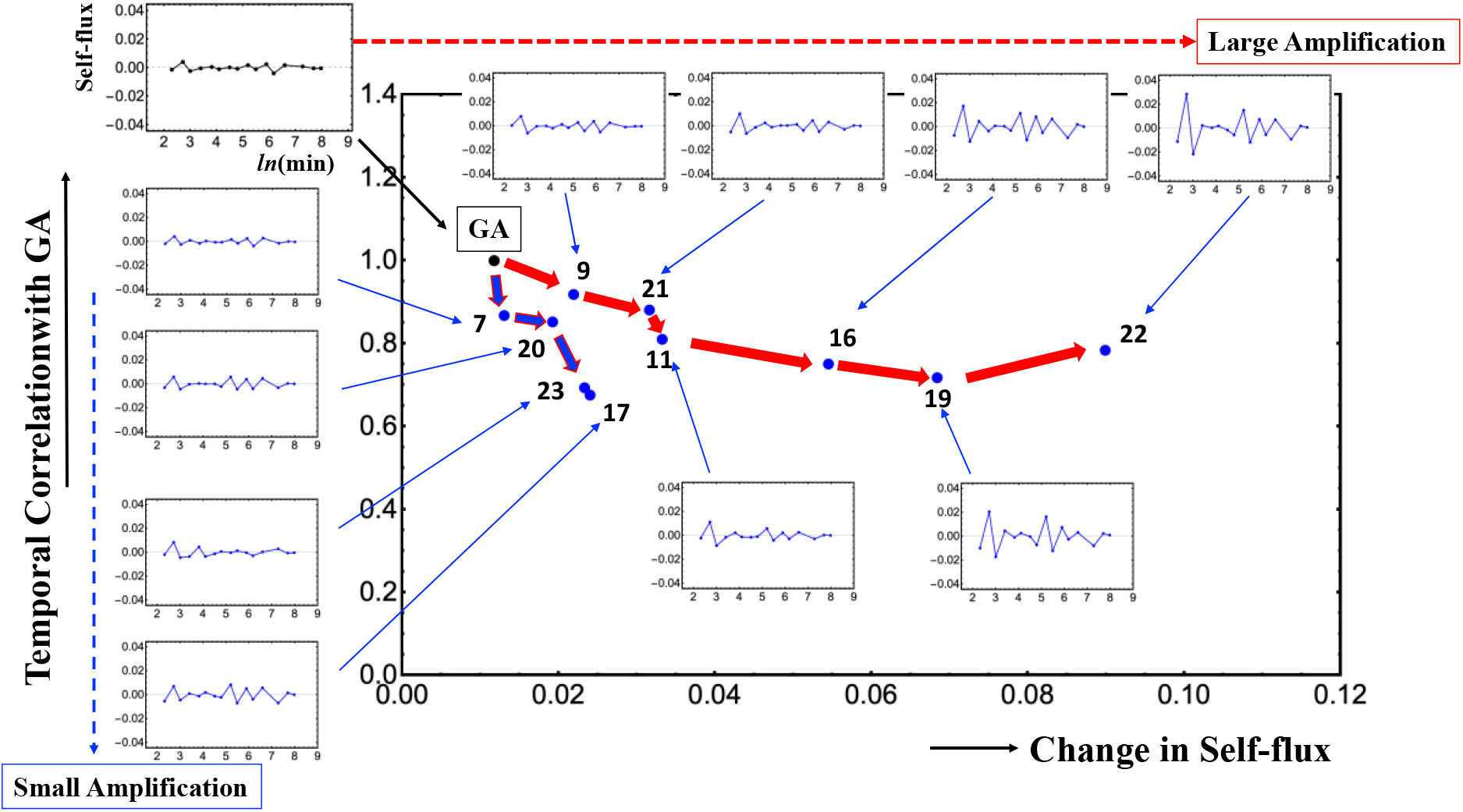
Amplification of temporal change in self-flux of the GA through the synchronized chromosomes. The GA is located at the edge of two paths (chromosomes 9 and 7), where the self-flux of the GA is amplified through them. This reveals temporal-spatio dynamics of the critical transition through the synchronized chromosomes. The *x*- and *y*-axes represents the same metrics as Figure 9.

Therefore, synchronization of the GA with the CP guides genome-wide avalanche to derive a super-critical genome state, where the synchronized wave spread through the synchronized chromosomes (Figure 10).

## 1.7 Discussions

The following discussions are updated ones as given in [Tsuchiya et al. 2016] with our recent further findings.

### 1.7.1 Biological Interpretations of the Global and Local Perturbations of SOC Control in MCF-7 Cells

EGF and HRG induced a transient and sustained dose-dependent phosphorylation of ErbB receptors, respectively, followed by similar transient and sustained activation kinetics of Akt and ERK [Nagashima et al. 2007]. After ERK and Akt activation from 5-10min, the ligand-oriented biphasic induction of immediate stress response key proteins of the AP-1 complex (c-FOS, c-JUN, and FRA-1) took place: high for HRG and negligible for EGF. The proteins of the AP-1 complex are non-specific stress-responders supported by the phosphorylation of ERK in a positive feedback loop. In addition, the key reprogramming transcription factor c-MYC (the protein of which peaked at 60 min, as confirmed in JE’s laboratory [Krigerts et al. 2021]) can amplify the transcription of thousands of active and initiated genes or direct-indirect targets and “open’ the chromatin by recruiting histone acetyltransferase [Frank et al. 2001)]. Saeki et al. [Saeki et al. 2009] revealed and Krigerts et al. [Krigerts et al. 2021] confirmed that after HRG this peak of the positive phase coincides with activation of FOSL1 suppressors FHL2, and DIPA, initiating the negative phase of stress-response genes abating their activity by 3h. This sustained positive phase followed by negative feedback allowed determination for differentiation to occur.

Thus, the continuity of the biological relay of the HRG-induced early (committing) and late transcription activities leads to a determination of differentiation from 3h (cell-fate change), which is necessarily coupled to the suppression of proliferation and stops the genome boost by the ERK pathway. Neither a sustained ERK dependent positive feed-back loop, nor the following negative feed-back is achieved in the case of EGF. Subsequently, these cells did not differentiate but continued proliferation. It is worth noting that the aforementioned genes have elevated temporal variability so further confirming the link between biologically and physically oriented interpretations of the proposed model [Zimatore et al. 2021].

In the corresponding expression data, we observed, after HRG, a powerful genome engine causing a pulse-like global perturbation (10-20min) as pre-committing and erasure of the initial-state CP to induce the genome-state change after 2h, which was not observed after treatment with EGF, in which case only local perturbation (i.e., only vivid activity of the super-critical state) was observed. The role of the constitutive pericentric domains PADs, mostly splitting into the low variance particles and losing their (H3K9me3+H3K4me3) barrier from H3K4me3-positive surrounding euchromatin within 5-10 min coinciding with start of this early CP genes response, perhaps allowing transmission the energy of this heterochromatin bursting to start the GA genome-engine, i.e., biological mechanism for the synchronization of dynamics of the CP with one of the GA – it should not be underestimated as this is a real physically registered process. As to observed in control the (H3K9me3+H3K4me3) “collar” around large PADs disappearing with their bursting HRG, it is interesting to refer to recent discovery of the same non-canonical bivalent chromatin form, shown as the partially methylated domain (PMD), which is protecting silent late replication timing [Du at al. 2021)].

Finally, it is important to note that the genome-engines (Figure 2) reveal that the sub-critical state (low-variance expression: a majority of RNA expressions) generates the global perturbation in the HRG response as well as in other biological regulation even in embryo development. The dynamic control of gene expression in the sub-critical state is expected from the cooperative ensemble behaviors of genomic DNA structural phase transitions through interaction with environmental small molecules, which has not been considered in previous biological studies. Thus, a true biological picture for cell-fate change may be obtained by deciphering the biological functions of genes in the sub-critical state in a coordinated (rather than an individual) manner.

### 1.7.2 Global Perturbations of SOC Control in HL-60 Cells Committed to Differentiation

Developmental biologists discriminate two modes of differentiation process: i) Reversible, with the capability of autonomous differentiation and ii) essentially irreversible [Barresi and Gilbert 2020]. The reaching of a ‘commitment’ state toward terminal differentiation ranges between 8 to 24h [Tarella et al. 1982; Tsiftsoglou et al. 1985] shorter than the duration of a single doubling cycle.

Consistent with these findings, we revealed global perturbation involving critical states to DMSO at 18h and to atRA at 24h [Tsuchiya at al 2020]. The achievement of cell-fate determination after 18h for DMSO and after 24h for atRA (Figures 3B,C) occurs in these two models in different ways, although they converge at the same final state at 48h.

In particular, we observed early global genome perturbation in the response to atRA (at 0-4h: see Fig. 16 in [Tsuchiya at al 2020]), which was not seen in the DMSO model. This may be due to a difference in Ca2+ influx [Papp et al. 2012] which controls key cell-fate processes including fertilization, early embryogenesis [Whitaker 2006], and differentiation [Yen et al. 1987; Schaefer et al. 1994; Dolmetsch et al. 1997; Launay at al. 1999].

### 1.7.3 SOC Control in Human and Mouse Related to the Developmental Oocyte-to-Embryo Transition

Fertilized mature oocytes change their state to become developing embryos. This process implies a global restructuring of gene expression. The transition period is dependent on the switch from the use of maternally prepared stable gene transcripts to the initiation of proper embryonic genome transcription activity. It has been firmly established that, in mice, the major embryo genome activation occurs at the two-cell stage (precisely between the mid and late 2-cell states), while in humans this change occurs between the 4- and 8-cell stages [Telford et al. (1990)].

These time intervals are detected by our SOC analysis. The switching of the genome-engine (Figures 4A,B) after the 4-cell state (at the 8-cell state) for human embryo development and after the late 2-cell state (at 4-cell state) for mouse embryo development. These timings coincide with those of the erasure of initial sandpile critical behaviors (see Fig.7 in [Tsuchiya at al. 2016]), where the CP is the summit of the sandpile critical behavior.

The erasure of paternal imprinting involves DNA 5-methylcytosine de-methylation and hydroxymethylation. This process allows fort de-compaction of the repressive heterochromatin and a consequent increase in the flexibility of the transcribing part of chromatin [J.S. Choy et al. 2010]. This process increases gradually and reaches a maximum 2-cell stage in mouse and 8-cell stage in human embryo [H. Guo et al. 2014; Niakan et al. 2012]. DNA de-methylation unpacks repressive heterochromatin, which manifests in the dispersal and spatio-temporal reorganization of pericentric heterochromatin as an important step in embryonic genome reprogramming [Probst et al. 2008; Yang et al. 2013].

In addition, transposable elements, which are usually nested and epigenetically silenced in the regions of hypermethylated constitutive heterochromatin, also become temporarily activated during the oocyte-to-embryo transition in early embryogenesis [Peaston et al. 2007; Guo et al. 2014] likely creating a necessary critical level of transcriptional noise as a thermodynamic prerequisite for the non-linear genome-expression transition using SOC.

### 1.7.4 Genome Computing

The CP (a set of critical genes) act as the organized center of cell-fate. The CP at critical transition exhibits bimodal domains [Tsuchiya et al. 2020a], which suggests folding and unfolding chromatin dynamics through synergistic or cooperative behavior in higher-order structural transition of genomic DNA (see Figure 1C). This is recently confirmed by the evidence of a massive re-arrangement of pericentromere-associated domains (PADs) [Krigerts et al. 2021]. The necessary link between chromatin remodeling and gene expression regulation was experimentally assessed [Wachsmuth et al. 2016]. This bi-phasic CP genes, thus, suggests the existence of a complex network of mega-sized ON/OFF DNA phase transitions in the cell-fate change.

Yoshikawa’s group [Yoshikawa and Matsuzawa 1996; Takagi and Yoshikawa 1999; Yoshikawa and Noguchi 1999] revealed that the time-dependent behavior of the genomic DNA transition follows a kinetic equation to exhibit cubic characteristics generated in the first-order phase transition. A simultaneous change in the translational and conformational entropy of giant DNA together with surrounding counter ions causes bimodality in the free energy, which in turn control the manner of folding transition as derived from functional derivative of the free energy. The kinetics represented with cubic nonlinearity corresponds to fundamental characteristic nerve firings [FitzHugh 1955; Izhikevich et al. 2006]. Note that cubic nonlinearity in the time-differentiation equation corresponds to a kinetic representation of the bimodal free energy and in general, criticality behavior is interpreted based on the symmetry argument under this type of energy landscape of first-order phase transition in general [Ginzburg and Landau 1965].

Therefore, the state change in the CP is expected to be a computing process through coordinated high-ordered DNA structural transitions (see Figure 10) for guiding the cell-fate change. The CP may further act as the center of genome computing, where chromatin remodeling is the material basis of such computation. Fundamental understanding of ‘genome computing’ of the CP could lead to a novel control mechanism of the cell-fate change in a desirable manner.

## 1.8 Conclusion: A Unified Genomic Mechanism

The biological systems appear as staying ‘on the edge of chaos’ oscillating around an equilibrium state without reaching it, being continuously challenged by the vagaries of their microenvironment. The continuous adaptation to these challenges so to keep a global homeostasis is the task of the so-called ‘hot spots’, genes having an elevated temporal variance mirroring such endless adaptive process. This temporal variance is both the source of long-term stability and, when reaches a given threshold, the determinant of cell-fate transition. Self-organized critical (SOC) control is responsible for the massive reprogramming of genome expression in cell-fate transition. Our integrated approach based on physics, statistics and biology provides further insights on a unified genomic mechanism of cell-fate change: how and when cell-fate change occurs on the genome [Tsuchiya et al. 2020a]. Key updated features can be summarized as:

1. An autonomous control genomic system develops the genome-engine in whole genome expression, which is coordinated by the emergence of a critical point (CP: a peculiar set of genes) at both the cell-population and single-cell levels. The CP act as the organized center of cell-fate, where the CP and GA (genome-attractor) form a phase segregation (bimodal domain) with free energy barrier. Depressing the free energy barrier, the switching of the genome-engine occurs through cell-fate change to induce coherent change in genome expression.
2. Possible biological mechanism of depressing free energy barrier (based on first-order phase-transition) might stem from spatial interaction of two chromatin types - silencing pericentric (hetero-) chromatin domains (PADs) and transcribing euchromatin.
3. The CP is activated by ‘hot-spots’ genes. When hot-spots oscillation exceeds a given threshold, the CP mediated critical transition can take place to induce change in the GA expression in a synchronized manner. This synchronized wave spread over synchronized chromosomes in the genome to amplify the change in the GA for global expression avalanche.
4. The switching oscillation between active and inactive chromatin compartments is the material basis of the hot-spots oscillation. Previous studies reported a large degree of spatial plasticity of the switching chromatin compartments across cell types, with approximately 36% of the genome compartments [Dixon et al. 2015]. This chromatin switching is the responsible of massive change in expression and is mirrored by fusion/splitting dynamics of PADs [Krigerts et al. 2021].

As for the development of a theoretical foundation for the autonomous SOC critical control mechanism as revealed in our findings is expected to open new doors for a general control mechanism of the cell-fate change and genome computing. The unveiling of genome structure-expression relationships is here to stay.

## 1.9 Methods

### 1.9.1 Biological Data Sets

Detail descriptions of seven distinct biological regulation data was provided in Methods [Tsuchiya et al. 2020a]. Here short descriptions are given with experimental time points. These data were considered examples of a general mechanism and have been selected based on the availability of a sufficient number of experimental time points.

#### Cell population

1. Heregulin (HRG)-stimulated MCF-7 human breast cancer cells compared with non-differentiated epidermal growth factor (EGF)-stimulated MCF-7 cells [Saeki et al. 2009]; 18 time points (Microarray data: GEO ID: GSE13009): *t*_1_ = 0, *t*_2_ = 10,15, 20, 30, 45, 60, 90min, 2, 3, 4, 6, 8, 12, 24, 36, 48, *t*_T = 18_ = 72h,
2. All-trans retinoic acid (atRA)- and dimethyl sulfoxide (DMSO)-stimulated HL-60 human leukemia cells differentiated to neutrophil cells [Huang et al. 2005]; 13 time points (Microarray data: GSE14500): *t*_1_= 0, *t*_2_ = 2, 4, 8, 12, 18, 24, 48, 72, 96, 120, 144, *t*_T=13_ =168h,

#### Single cell

3. Differentiation of T helper 17 (Th17) cells from naïve T helper (Th0) cells [Ciofani et al. 2012]: 9 time points (RNA-Seq data: GSE40918): *t*_1_ = 0, *t*_2_ = 1, 3, 6, 9, 12, 16, 24, *t*_T=9_ = 48h.
4. Mouse early embryo development [Deng et al. 2014]: 10 cell states (RNA-Seq data: GSE45719): zygote, early 2-cell, middle 2-cell, late 2-cell, 4-cell, 8-cell, morula, early blastocyst, middle blastocyst and late blastocyst,
5. Human early embryo development [Yan et al. 2013]: 7 cell states (RNA-Seq data: GSE36552): Human: oocyte, zygote, 2-cell, 4-cell, 8-cell, morula and blastocyst.

Note: For microarray data, the Robust Multichip Average (RMA) was used to normalize expression data for further background adjustment and to reduce false positives [Bolstad et al. 2003; Irizarry et al. 2003; McClintick and Edenberg 2006]. For RNA-Seq data, RNAs with RPKM values of zero over all of the cell states were excluded.

### 1.9.2 Normalized Root Mean Square Fluctuation (*nrmsf*)

*Nrmsf* (see more Methods in [Tsuchiya et al. 2015]) is defined by dividing *rmsf* (root mean square fluctuation) by the maximum of overall {*rmsf*_i_}:

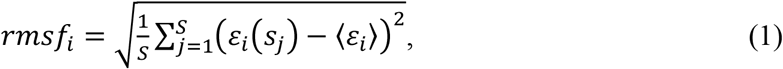

where *rmsf*_*i*_ is the *rmsf* value of the *i*^*th*^ RNA expression, which is expressed as *ε*_*i*_(*s*_*j*_) at a specific cell state *s*_*j*_ or experimental time (e.g., in mouse embryo development, *S* = 10: *s*_1_ *=* zygote, early 2-cell, middle 2-cell, late 2-cell, 4-cell, 8-cell, morula, early blastocyst, middle blastocyst and *s*_10_ *=* late blastocyst), and 〈*ε*_*i*_〉 is its expression average over the number of cell states. Note: *nrmsf* is a time-independent variable.

### 1.9.3 Updated Expression Flux Analysis

Here, we update the expression flux analysis to describe the genome-engine mechanism on both single cell and population genome expression (refer to Methods in [Tsuchiya at al. 2020a]). The dynamics of coherent behavior emerges from stochastic expression in distinct critical states (coherent-stochastic behavior: CSB [Tsuchiya et al. 2015]), which follows the dynamics of the CM of a critical state as a critical-state attractor. A *heteroclinic critical transition* guides coherent behaviors emerged in the (super-, near- and sub-critical) critical states. The genome-engine represents overall picture of time-average between local critical-state attractors, which corresponds to a stable manifold of the thermodynamically open system. When cell-fate change occurs, temporal perturbation on the genome-engine coherently changes and switches through critical transition (Figures 3,4). At the switching, all fluctuations from the genome-engine pass zero: the genome system itself lies in a stable point (*non-equilibrium fixed point*) of thermodynamically open system.

Essential points of the update are as follows:

1. Instead of actual experimental time, the meaning of ‘a time’ in whole gene expression represents ‘a state’ of cells or single cell, such that experimental times can be represented just by specific experimental events (whole numbers) with the following two biophysical reasons:

a. Temporal expression variance, *nrmsf* acts as the order parameter for the self-organization. Once whole expression at *t* is ordered and grouped (*k* groups) by *nrmsf,* sandpile-type transitional behavior (Figure 1) emerges in the change in group expression between different times, which shows that time is an implicit parameter in SOC. The top of the sandpile is a critical point (CP), where the CP usually exists at around a zero-fold change (null change in expression). Cell-fate change occurs after erasure of initial-state CP, where the sandpile-type profile between *t*_0_ and *t*_c_ disappears, i.e., cell-fate change occurs implicitly at *t*_c_.
b. In embryo development, whole numbers correspond to cell-states such as zygote and 2-cell states. Therefore, to develop a unified expression flux analysis, times and cell-states are represented by whole numbers
2. Representation of the self-flux is updated to clearly show a formation of cyclic state-flux between-critical states (see Figure 2). The numerical value of the *i*^th^ critical state, *X*_*i*_(*t*_*j*_) (i.e., *i* = super-, near- or sub-critical state) is represented by |*i*⟩(*t*_*j*_) at a specific experimental event (*t*_j_). The net self-flux, ⟨*i*|*i*⟩(*t*_*j*_) describes the effective force on CM of the *i*^th^ critical state, which represents the difference between the positive sign for incoming force (net IN self-flux) and the negative sign for outgoing force (net OUT self-flux):

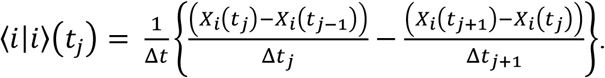 Interaction flux from the *j*^th^ to *i*^th^ crucial state is defined as

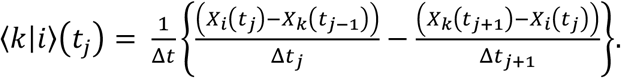 Then, the self-flux, ⟨*i*|*i*⟩(*t*_*j*_) can be expanded by interaction and external flux (see details in [Tsuchiya at al. 2020b]):

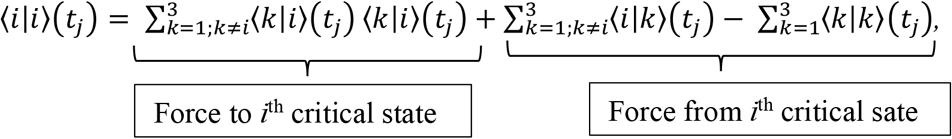

where the first term describes the force from other critical states to the *i*^th^ critical state, i.e., internal force within the heteroclinic system, the second and third terms represent the force from the *i*^th^ critical state force to other critical states, and momentum change in cell nucleus (environment), 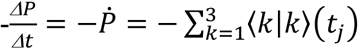, respectively. As an example, the self-flux of 1^st^ critical state (super-critical state), 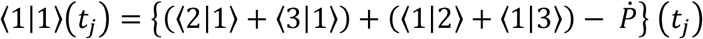. Note that

1. 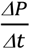 term describes momentum change in the heteroclinic system, i.e., force coming from cell environment, and
2. The second and third terms are previously considered as a single term describing incoming flux (positive sign) from the environment to a critical state or outgoing flux (negative) from a critical state to the environment.

## Acknowledgments

MT sincerely thanks the following institution and individuals who helped complete this research project: the SEIKO Life Science Laboratory, Osaka, Japan and his family (particularly, his daughters: Drs. Kimiko and Kazumi Tsuchiya, and Dr. Harry Taylor with any editing).

## Funding

This study was supported in part by Japan Society for the Promotion of Science; JSPS KAKENHI Grant Number JP20H01877 and by European Regional Development Fund; Project Number: 1.1.1.1/18/A/099.

## Notes

The authors declare no conflict of interest.

### Competing Interest Statement

The authors have declared no competing interest.

